# Defining the order of assembly of the *Clostridioides difficile* divisome complex

**DOI:** 10.64898/2026.01.19.700290

**Authors:** Gregory A. Harrison, Pola Kuhn, Shailab Shrestha, Paula Caballero Blanco, Larissa Havey, Aimee Shen

## Abstract

Cell division is the ancient pathway by which bacteria synthesize a septum of peptidoglycan, dividing the cell into two. Where all walled bacteria were previously thought to use FtsW-FtsI orthologs to synthesize septal peptidoglycan during division, we recently discovered that the major pathogen *Clostridioides difficile* is missing FtsW-FtsI and instead relies on the activity of the bifunctional Class A PBP called PBP1 to synthesize the septal peptidoglycan. Furthermore, *C. difficile* either does not encode or require the majority of canonical divisome proteins described in model bacteria aside from the divisome protein orthologs FtsZ, SepF, and ZapA. Indeed, unlike model systems, SepF and ZapA are essential in *C. difficile*, suggesting that they have evolved to have a critical function in cell division without the redundant mechanisms present in model organisms. Thus, *C. difficile* uses a fundamentally different division mechanism compared to previously studied bacteria. To understand how this unusual complex is assembled in *C. difficile*, we combine CRISPR interference (CRISPRi)-based knock-downs with fluorescent fusions to determine that the hierarchical order of assembly occurs in three phases: (i) FtsZ/ZapA, (ii) SepF, and (iii) PBP1. We further investigate the order of assembly of several non-essential mid-cell localizing proteins and discover that MldA, MldC, DivIVA, FtsK, and PBP3 depend on FtsZ, SepF, and PBP1 for localization, whereas MldB localizes independently of SepF and PBP1. Our work provides a model for divisome assembly in *C. difficile* and validates genetic and cytological tools that can be used to mechanistically dissect this pathway in the future.

**IMPORTANCE:** Bacterial cell division has been extensively studied in model systems, but little is known about how this essential process occurs in the clinically important pathogen *Clostridioides difficile*. Studies in model systems have shown that cell division is carried out by a large multi-protein complex called the “divisome.” While components of the divisome are widely conserved and can be traced back to the last bacterial common ancestor billions of years ago, *C. difficile* uses a unique mechanism of division that is independent of the majority of these canonical divisome genes. In the current study, we characterize the core, essential divisome comprised of FtsZ, ZapA, SepF, and PBP1, and build a model for the order of assembly of this unusual divisome complex.

## INTRODUCTION

The essential process of bacterial cell division is driven by a large multi-protein complex known as the divisome. Divisome assembly centers around the tubulin-like protein FtsZ, which assembles into a ring at the mid-cell and serves as a scaffold to organize the divisome complex (1–3). Divisome assembly is a multi-step process that broadly occurs in three stages: (i) FtsZ and other early divisome proteins mark the site of division, (ii) transmembrane proteins are recruited, including the septal peptidoglycan (PG) synthases, and (iii) when assembly is complete, the PG synthases are activated, leading to septum synthesis, which drives cytokinesis.

In the model rod-shaped Firmicute species *Bacillus subtilis*, the localization of FtsZ is coordinated by multiple divisome-associated proteins that promote bundling of FtsZ filaments and/or tethering of the filaments to the membrane, including ZapA, SepF, FtsA, EzrA, and GpsB (4–14). These non-essential proteins have partially overlapping functions to promote FtsZ-ring assembly and/or membrane anchoring and are therefore each individually dispensable; however, they become synthetically lethal in higher order mutant combinations (4, 6–9, 11, 12, 15, 16).

In *B. subtilis* and other model systems, proper assembly of the FtsZ proto-ring in the cytosol allows this cytoskeletal scaffold to recruit trans-envelope proteins, including the regulatory sub-complex comprised of FtsQ (DivIB), FtsL, and FtsB (DivIC) (17–22). The FtsQ-FtsL-FtsB sub-complex promotes the recruitment and activation of the FtsW-FtsI septal peptidoglycan synthase complex at the site of division (23–36). Each of these proteins is essential for division in standard laboratory conditions in *B. subtilis* (18, 20, 37, 38), and the genes encoding FtsQ, FtsL, FtsW, and FtsI have been traced to the last bacterial common ancestor billions of years ago (39). Indeed, the FtsW-FtsI synthase complex was previously considered essential for septum synthesis in virtually all bacteria with a cell wall (3, 23, 39, 40).

Despite the extreme conservation of divisome components across most bacteria (39), we recently showed that *C. difficile* exhibits marked differences in the composition of its divisome relative to *B. subtilis* and previously studied bacteria. Specifically, we found that *C. difficile* does not encode orthologs of FtsW and FtsI for mediating vegetative cell division (41). Additionally, the genes encoding the typically essential divisome regulators FtsQ, FtsL, and FtsB are completely dispensable for vegetative cell division and have evolved to function specifically during sporulation (41). In lieu of FtsW-FtsI, we showed that *C. difficile* uses its sole class A penicillin binding protein (aPBP), PBP1, to drive the synthesis of the septum during vegetative cell division (41). The essential function for an aPBP during vegetative cell division contrasts with previously studied bacteria, where aPBP enzymes are important for fortifying the cell wall (42–44), filling in gaps in the peptidoglycan mesh (45–48), reinforcing the septum synthesized by FtsW-FtsI (49–51), or driving cell elongation at the poles (52–57), depending on the species. Thus, the essential role of PBP1 in synthesizing vegetative division septa in *C. difficile* represents to our knowledge the first example of an aPBP driving cell division in the absence of the canonical FtsW-FtsI synthase complex.

Intriguingly, *C. difficile* encodes orthologs of ZapA and SepF, but it lacks identifiable orthologs of the FtsA, EzrA, or GpsB divisome proteins found in the Firmicutes, suggesting that regulation of FtsZ in *C. difficile* differs substantially from *B. subtilis*. Since the majority of the expected divisome proteins in *C. difficile* are either missing from the genome (FtsA, EzrA, GpsB, FtsW, FtsI), or are dispensable for division (FtsQ, FtsL, FtsB), the molecular details of how *C. difficile* carries out cell division remain largely unknown.

Recent work has shown that at least 24 different proteins localize to mid-cell in *C. difficile* (41, 58–65), of which 8 were predicted to be essential by transposon-insertion sequencing (63, 66). Four of these proteins are likely essential for reasons unrelated to cell division (RodA, PBP2, MreC, and MreD), although it is possible they play an auxiliary function in the division apparatus (41, 63). Therefore, of the essential, mid-cell localizing proteins, we are primarily interested in the predicted proto-ring components, FtsZ, ZapA, and SepF, and the primary septum synthase, PBP1.

While FtsZ has been localized to mid-cell and is essential for division in *C. difficile* (60, 67), its regulation remains poorly understood. *C. difficile* ZapA also localizes to mid-cell and is essential for division (61, 63), although this essentiality appears to be unique to *C. difficile* because it is dispensable in most previously studied bacteria. While ZapA has been shown to promote FtsZ filament bundling and FtsZ-ring stability by mediating lateral interactions between filaments in model systems (4, 68–70), it is likely non-essential due to the presence of alternative mechanisms to bundle FtsZ filaments and promote FtsZ-ring stability (4, 71–77). While these redundant mechanisms presumably do not exist in *C. difficile*, the precise mechanism by which ZapA regulates the divisome complex has not yet been examined.

In addition to ZapA, the *C. difficile* SepF ortholog localizes to mid-cell (63) and is predicted to be essential (63, 66), but its requirement for cell division in *C. difficile* has not yet been examined. In model systems, SepF interacts with FtsZ and associates with the membrane, serving as an FtsZ-membrane anchoring protein (5, 6, 11). However, in Firmicutes species that encode other membrane anchors such as FtsA, SepF is not required for stable FtsZ-ring assembly, although SepF-deficient cells exhibit septal irregularities (5, 78). Interestingly, SepF is considered the ancestral FtsZ membrane anchor, as bacterial and archaeal species that lack other membrane anchors, such as FtsA, often encode a SepF ortholog (79); for example, it is essential for FtsZ-ring assembly in Actinobacteria species that lack FtsA (80–82). Since SepF is surprisingly not required for stable FtsZ-ring assembly in the archeal species in which it has been studied, additional unknown mechanisms for stabilizing and tethering the FtsZ ring likely exist in those Archaea (83).

In this study, we use CRISPR-interference (CRISPRi) and fluorescent fusion proteins to FtsZ, ZapA, SepF, and PBP1 to investigate the assembly of the *C. difficile* divisome. Using CRISPRi-compatible trans-complementation, we develop genetic tools to assess the function of fluorescently-tagged divisome proteins. Additionally, we investigate the order of assembly of *C. difficile*’s core essential divisome proteins and examine the role of several non-essential auxiliary cell division proteins in this pathway. These studies provide insight into the non-canonical divisome used by *C. difficile* to mediate vegetative cell division and establish a framework for future studies of this important process.

## RESULTS

### SepF is essential for septum synthesis in *C. difficile*

Based on Tn-seq data, there are four genes encoding known or predicted divisome proteins that are predicted to be essential in *C. difficile*: *ftsZ*, *zapA*, *sepF*, and *pbp1* (63, 66). Prior work using CRISPRi-based knock-down (KD) showed that *ftsZ*-KD, *zapA*-KD, and *pbp1*-KD each result in cells with a long, filamentous morphology (41, 63, 67), which is a hallmark phenotype of cells that cannot divide (84–87). Since *ftsZ*, *zapA*, and *pbp1* are each either transcribed as a sole gene or are the last gene in their operon, based on RNA sequencing and transcription start site mapping (88), the filamentation phenotypes can be attributed to the specific knock-down of these genes. However, because the *sepF* gene is embedded within an operon that contains *ylmD*, *ylmE*, *sepF*, *ylmG*, *ylmH*, and *divIVA* (**Fig. 1A**), the role of *sepF* in *C. difficile* division has yet to be directly addressed thus far. While prior work that CRISPRi-KD of the entire *sepF*-containing operon using a sgRNA targeting *ylmD* induced cell filamentation (63), the specific gene(s) in this operon that are required for cell division remain unclear.

**Fig 1:**
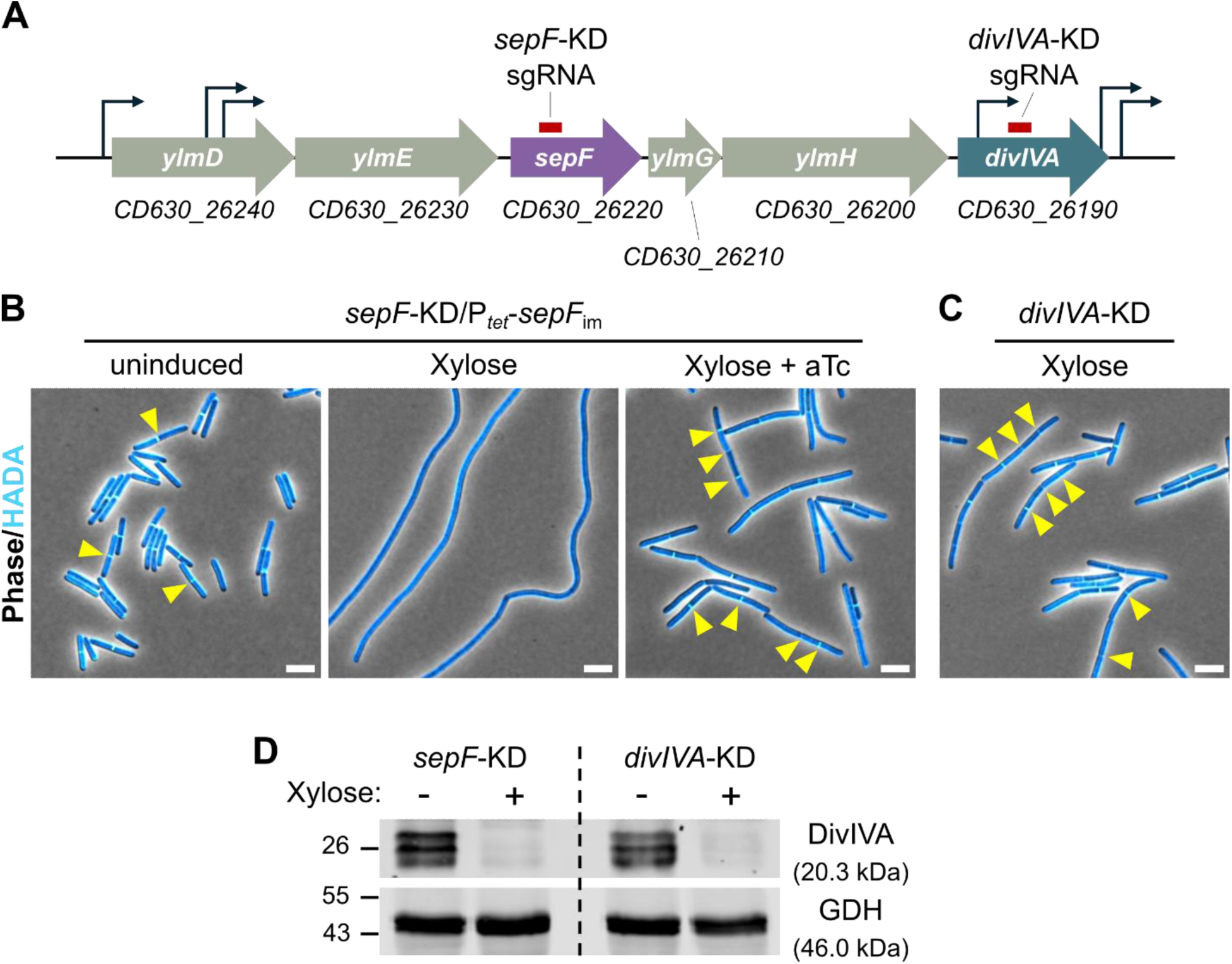
SepF is required for the synthesis of division septa. (A) Organization of the operon containing *sepF*. The location targeted by CRISPRi sgRNAs are indicated. Both *sepF* and *divIVA* sgRNAs target the non-template strand. (B) A *C. difficile* strain was generated harboring a plasmid-encoded xylose-inducible CRISPRi *sepF*-KD cassette in addition to a CRISPRi-resistant *sepF*_im_ complementation cassette integrated in the genome under control of an aTc-inducible P*_tet_* promoter. Culturing in 2.5% xylose results in induction of the *sepF*-KD cassette, and addition of 2.5 ng/mL aTc induces expression of the *sepF*_im_ complementation construct. Cells were labeled with HADA to visualize sites of peptidoglycan synthesis and/or remodeling, then fixed for microscopy. Yellow arrows indicate division septa. Scale bars = 5 μm. (C) A *C. difficile* strain harboring a plasmid-encoded xylose-inducible CRISPRi *divIVA*-KD cassette was cultured in the presence of 2.5% xylose, then labeled with HADA and fixed for microscopy. Yellow arrows indicate division septa. Scale bars = 5 μm. (D) *C. difficile* harboring a plasmid-encoded xylose-inducible *sepF*-KD or *divIVA*-KD CRISPRi cassette were cultured in the absence (-) or presence (+) of 2.5% xylose for approximately 6 hr (∼8 doublings) and western blotting was performed for DivIVA with GDH as a loading control. Data are representative of at least two independent experiments.

While *sepF*, *ylmG*, and *divIVA* all encode proteins known to localize to the site of division (63, 65), we focused on the potential role of *sepF* due to its known role in division in other species (5–7, 79–81, 83, 89). To directly test if *sepF* is required for cell division, we used a xylose-inducible CRISPRi system targeting *sepF* combined with CRISPRi-compatible complementation (62, 90, 91). We have used this type of approach to complement CRISPRi-KD of essential genes in *C. difficile* (62), and similar approaches have been used in *Borrelia burgdorferi* and *Mycobacterium tuberculosis* (90, 91).

CRISPRi-KD of *sepF* caused a block in cell division, resulting in long filamentous cells defective in septum formation, as indicated by an absence of HADA incorporation into septa within the filaments (**Fig. 1B**). Since repression of *sepF* is also expected to knock-down expression of *ylmG*, *ylmH*, and *divIVA* based on the predicted operon structure, we then tested if complementing with *sepF* alone could restore septum synthesis in the KD strain. To do this, we integrated a *sepF* complementation construct into the genome under the control of an anhydrotetracycline (aTc)-inducible P*_tet_* promoter (92) containing synonymous point mutations in the sgRNA-targeted sequence, rendering the construct “immune” to CRISPRi-targeting (*sepF*_im_). We then titrated the aTc concentration to identify the lowest concentration of aTc that enabled complementation of the KD. At 2.5 ng/mL aTc, induction of *sepF*_im_ restored septum synthesis in the *sepF*-KD strain (**Fig. 1B**). However, complemented cells developed a chaining phenotype. Therefore, while our data demonstrates that *sepF* is specifically essential for septum synthesis, cell chaining in the *sepF*-KD/*sepF*_im_ complementation strain suggests that decreased *ylmG*, *ylmH*, or *divIVA* expression likely prevents efficient cell separation.

Chaining has been reported in mutant strains of *Streptococcus pneumoniae*, *S. suis*, and *Listeria monocytogenes* lacking *divIVA* (93–97). We therefore examined whether knocking down *divIVA* is sufficient to induce cell chaining using CRISPRi. We found that *divIVA*-KD phenocopied the cell chaining in the *sepF*-KD/*sepF*_im_ complemented strain (**Fig 1C**), suggesting that the chaining phenotype in this strain is likely caused by a lack of *divIVA* expression. Consistent with this conclusion, we found that DivIVA protein levels are decreased in the *sepF*-KD similar to the *divIVA*-KD (**Fig 1D**), verifying that *sepF*-KD strain is depleted of DivIVA due to polar effects on the operon. Together, our findings strongly suggest that SepF is critical for septum synthesis, while reduced DivIVA levels in the complementation of *sepF*-KD with *sepF*_im_ results in a secondary cell chaining phenotype.

### The FtsZ-ZapA-SepF proto-ring assembles at mid-cell earlier than PBP1

To compare the localization of these proteins to the *C. difficile* divisome, we integrated the expression constructs into the genome under control of an aTc-inducible promoter and titrated the aTc concentration to identify a level of fusion protein production that recapitulates their septal localization without inducing morphological abnormalities. We found that low-level expression of *ftsZ*-*mScI3*, *mScI3*-*zapA*, *sepF*-*mScI3*, and *mScI3*-*pbp1* allowed all the fusion proteins to localize to mid-cell (**Fig. 2A**), similar to prior studies (41, 60–63). When we quantified the enrichment of the proteins at the mid-cell relative to the sidewall or cytosol, we found that FtsZ, ZapA, and SepF fusions were highly enriched (≥5-fold) at the mid-cell, whereas PBP1 exhibited a modest mid-cell enrichment of ∼1.8-fold (**Fig S1**). Transmembrane proteins that are enriched at mid-cell are expected to be enriched by more than 2-fold above the sidewall, as the mid-cell will have two membranes once the septa are complete. Thus, our data suggests that PBP1 is localized throughout the cell, which may be consistent with PBP1 having a role in both the synthesis of the septum and the sidewall (41, 67). Additionally, by western blot analysis, we found that there is some level of cleavage of the mScI3 fluorescent fusion from each of these proteins (**Fig S2**), which likely also confounds our ability to precisely quantify protein localization within the cell. With these minor limitations in mind, we used these fluorescent fusions to learn about the relative order of assembly of the divisome complex in *C. difficile*.

**Fig 2:**
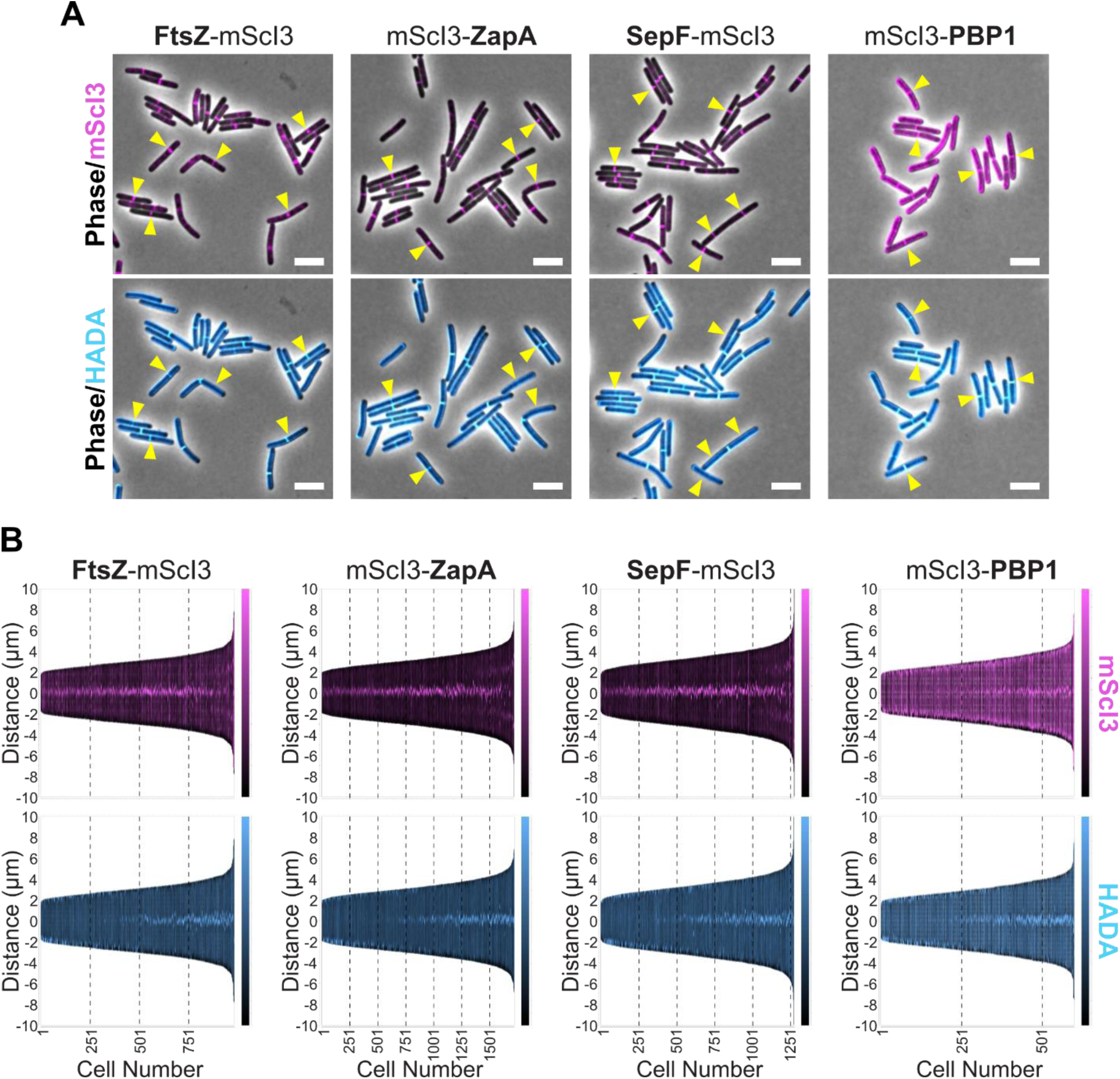
Localization profile of essential *C. difficile* divisome proteins. (A-B) *C. difficile* strains were made harboring expression cassettes to produce mScarlet-I3 (mScI3) fusion proteins under control of an aTc-inducible P*_tet_* promoter from an ectopic site in the genome. To induce production of the fusion protein, logarithmically growing cells were cultured in the presence of the following concentrations of aTc for 1 hr: 0.5 ng/mL aTc for FtsZ-mScI3 and mScI3-ZapA, 1 ng/mL aTc for SepF-mScI3, and 2.5 ng/mL aTc for mScI3-PBP1. Cells were labeled with HADA for 10 min to visualize sites of peptidoglycan synthesis and/or remodeling and then fixed for fluorescence microscopy. (A) Merged images containing phase and mScI3 signal (top) or phase and HADA signal (bottom) are shown, and yellow arrows point to examples of divisome protein foci. Scale bars = 5 μm. (B) Demographs were generated using MicrobeJ to visualize the medial axis fluorescence profile of ≥500 cells. Cells are ordered by length, and the mScI3 or HADA signal along the distance of the cell relative to mid-cell is shown in magenta and cyan, respectively. Data in A-B are representative of at least three independent experiments.

We used the MicrobeJ plugin in FIJI to generate demographs, which sorts cells based on their length, to visualize the medial fluorescence profile of each fluorescently-tagged protein across hundreds of cells. This allowed us to visualize the mid-cell mScI3 signal in cells at various stages of division. By monitoring HADA incorporation at mid-cell, it is possible to estimate whether the fluorescently-tagged protein localizes to the mid-cell prior to or concurrently with septum synthesis (98). The FtsZ-mScI3, mScI3-ZapA, and SepF-mScI3 fusions all localized to the mid-cell prior to the onset of septum synthesis (**Fig. 2B**), consistent with these proteins being early components of the divisome complex. In contrast, mScI3-PBP1 localized to mid-cell coincident with septum synthesis, as we reported in our prior work (62). These findings are consistent with FtsZ, ZapA, and SepF comprising the early proto-ring of the *C. difficile* divisome that assembles prior to recruitment of the trans-envelope proteins including PBP1 that ultimately trigger septum synthesis.

### Assessing the functionality of fluorescent protein fusions to essential divisome proteins

Although our fluorescent protein fusion constructs suggest that these divisome proteins localize to mid-cell in *C. difficile* (**Fig 2A**), similar to prior work (41, 60–63), it was unclear whether these fluorescent protein fusions are functional. Assessing the functionality of these tagged proteins is important for understanding whether there are limitations to using these fusions should they be found to be non-functional.

To determine the functionality of FtsZ-mScI3, mScI3-ZapA, SepF-mScI3, and mScI3-PBP1 fluorescent fusions, we used the CRISPRi trans-complementation system described above. Specifically, we combined xylose-inducible CRISPRi-KD cassettes with aTc inducible CRISPRi-“immune” complementation constructs encoding the fluorescent protein fusions. Each fusion carries a (GGGGS)_3_ linker between the mScI3 and protein of interest. While complementation of *ftsZ*-KD with the WT control *ftsZ*_im_ construct reversed the filamentation phenotype caused by *ftsZ*-KD (**Fig. 3A-B**), complementation with *ftsZ*_im_-*mScI3* did not, indicating that the FtsZ-mScI3 fusion is not functional (**Fig. 3B**). Notably, FtsZ-fluorescent protein fusions have often been found to be either non-functional (99) or only partially functional, with low temperatures being required for their proper function (8) in many model systems, so the localization of these fusions is typically analyzed in a merodiploid background.

**Fig 3:**
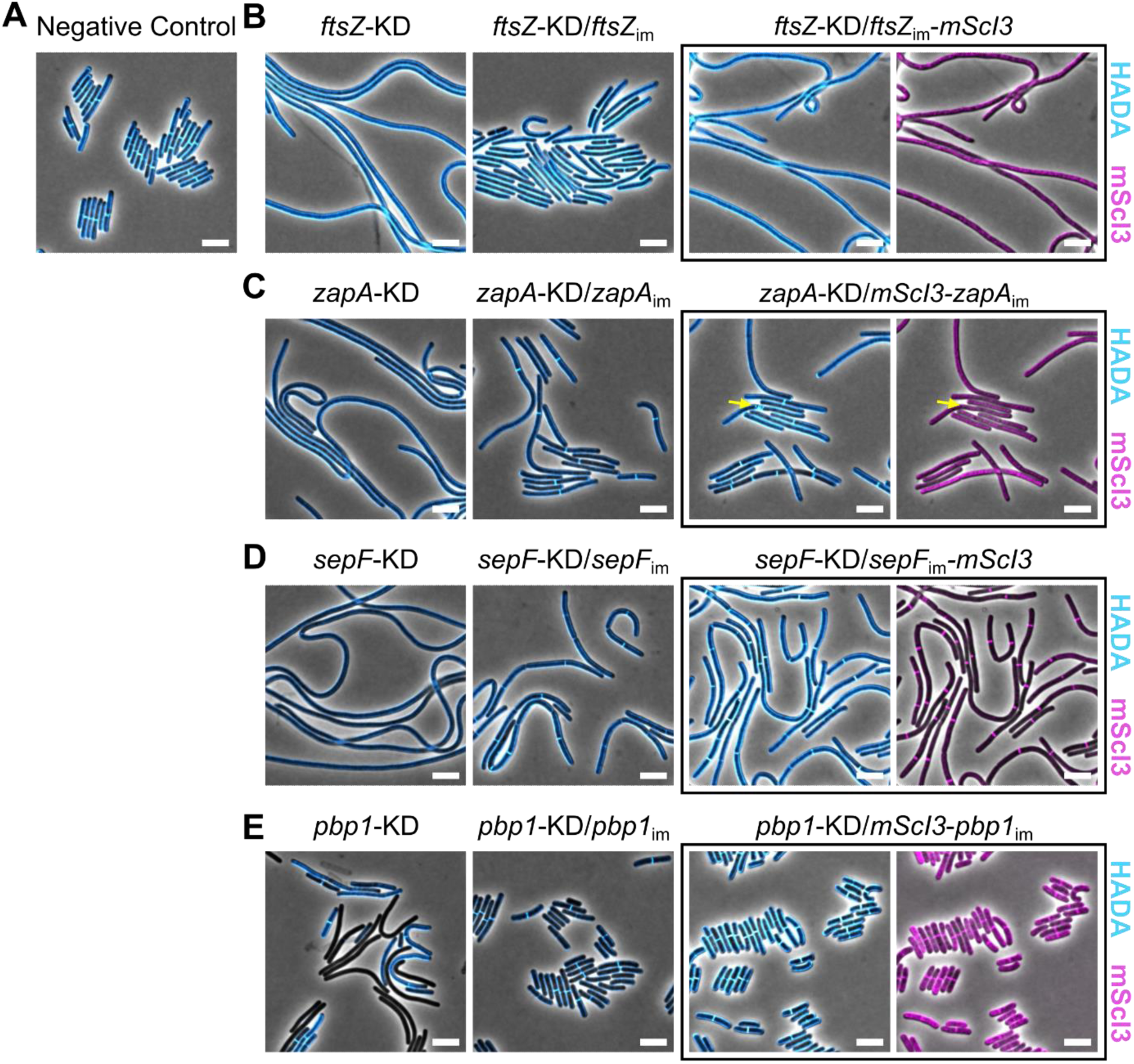
Assessing the functionality of fluorescent protein fusions to essential divisome proteins. (A-E) Plasmid-encoded xylose-inducible CRISPRi-KD cassettes that either (A) are non-targeting (negative control), or target (B) *ftsZ*, (C) *zapA*, (D) *sepF*, or (E) *pbp1*, were introduced into *C. difficile* strains carrying chromosomally-encoded complementation constructs that are “immune” to CRISPRi targeting. These CRISPRi-immune constructs were expressed under control of an aTc-inducible P*_tet_* promoter. The indicated strains of *C. difficile* were cultured for ∼6 hr in the presence of 2.5% xylose, to induce the CRISPRi-KD cassette, and aTc (*ftsZ*/negative control = 1 ng/mL; *zapA* = 50 ng/mL; *sepF* = 2.5 ng/mL; *pbp1* = 2 ng/mL), to induce expression of the CRISPRi-immune complementation construct. The cultures were then labeled with HADA to visualize sites of peptidoglycan synthesis and/or remodeling and fixed for microscopy. Merged images containing phase/HADA (blue) or phase/mScI3 (magenta) are shown. The HADA signal is normalized across all images, but the mScI3 signal was scaled such that the localization for each individual fusion protein was more easily detected. The yellow arrow in C points to an example of a HADA “spiral”; these structures were only observed in *zapA*-KD/*mScI3*-*zapA*_im_ cells. More examples of these structures can be found in Fig S3. All data are representative of at least two independent experiments. Scale bars = 5 μm.

In contrast, expression of *mScI3*-*zapA*_im_ rescued synthesis of septa and partially rescued the filamentation phenotype caused by *zapA*-KD (**Fig. 3C**). Since complementation with the WT *zapA*_im_ only partially rescued the filamentation phenotype, our ectopic *zapA* complementation cassette may not perfectly match the expression levels and/or regulation found at the endogenous gene locus. Notably, the mScI3-ZapA did not form distinct foci in the cells, perhaps due to over-expression of the fusion, masking the discrete mid-cell localization. We also observed the occasional formation of unusual spiral-like septa with the HADA label during conditional expression of *mScI3*-*zapA*_im_ (**Fig. 3C**, yellow arrow)(Fig. S3). Thus, while our data indicate that the mScI3-ZapA fusion can partially restore septum synthesis, it can also cause abnormal septum synthesis, perhaps due to incomplete bundling of the FtsZ-ring that directs the septum synthesis machinery. Regardless, these data indicate that mScI3-ZapA localization to mid-cell must be studied in a merodiploid background.

Notably, conditional expression of *sepF*_im_-*mScI3* in the *sepF*-KD strain enabled synthesis of division septa (**Fig. 3D**), similar to the *sepF*-KD/*sepF*_im_ complementation mutant (**Fig. 3D**)(**Fig. 1B**), although neither *sepF*_im_ construct was able to rescue the chaining phenotype that is presumably caused by the polar effects on *divIVA* expression. Therefore, these data indicate that SepF-mScI3 is at least partially functional. Finally, conditional expression of *mScI3-pbp1*_im_ also restored normal septum synthesis in the *pbp1*-KD strain (**Fig. 3E**), strongly suggesting that the mScI3-PBP1 protein fusion is also functional. Nevertheless, since fluorescent protein fusions to ZapA, SepF, or PBP1 undergo low levels of cleavage of mScI3 from the fusions, it is possible that the small amount of untagged protein produced is responsible for driving the complementation phenotype (**Fig S2**). While ruling out this possibility will require additional work to identify fluorescent fusions that are not cleaved, our data thus far suggest that mScI3-ZapA, SepF-mScI3, and mScI3-PBP1 are partially or fully functional in *C. difficile*, whereas FtsZ-mScI3 is clearly not functional.

### Hierarchical recruitment of *C. difficile* divisome proteins

We next sought to determine the order of assembly of these essential divisome proteins because in many systems the recruitment of the divisome proteins is hierarchical, where the recruitment of each subsequent protein to the complex is dependent on the proper assembly of the earlier proteins (22, 100). To examine the order of assembly of *C. difficile*’s core divisome complex, we analyzed the localization of the fluorescent fusion proteins in which *ftsZ*, *zapA*, *sepF*, or *pbp1* expression was knocked-down using CRISPRi. While our analyses indicated that ZapA, SepF, and PBP1 fluorescent fusions are at least partially functional (**Fig 3**), we nevertheless decided to visualize these fusions in the context of a fully functional divisome complex by employing the commonly-used dilute-label approach, in which the localization of fluorescently-tagged proteins is studied in a merodiploid background (101, 102).

When we knocked-down *ftsZ* expression, the mScI3-ZapA, SepF-mScI3, and mScI3-PBP1 fusions all failed to assemble into distinct foci, unlike in control cells (**Fig. 4A-B**). Therefore, the assembly of ZapA, SepF, and PBP1 at mid-cell depends on the presence of FtsZ, similar to previously studied model systems (4, 5, 103). When *zapA* expression was knocked-down, FtsZ-mScI3 and mScI3-PBP1 also failed to localize to distinct foci, suggesting that ZapA is critical for FtsZ and PBP1 localization (**Fig. 4B**). This is somewhat surprising, as ZapA in well-studied model systems is dispensable for division due to the presence of redundant mechanisms for promoting FtsZ-ring stability (4, 71–77). Thus, ZapA appears to have evolved a uniquely critical function in driving assembly of the divisome complex in *C. difficile*. This unique function likely explains why this gene is essential in *C. difficile* despite being dispensable in previously studied bacterial systems. Intriguingly, we observed that SepF-mScI3 localized to distinct foci in *zapA*-KD cells (**Fig. 4B**). Since the SepF-mScI3 puncta were irregularly spaced and less well defined than the crisp foci observed in control cells (**Fig. 4A**), SepF appears to only partially depend upon ZapA to assemble into foci at mid-cell, while FtsZ and ZapA are co-dependent for assembly into foci (**Fig. 4C**).

**Fig 4:**
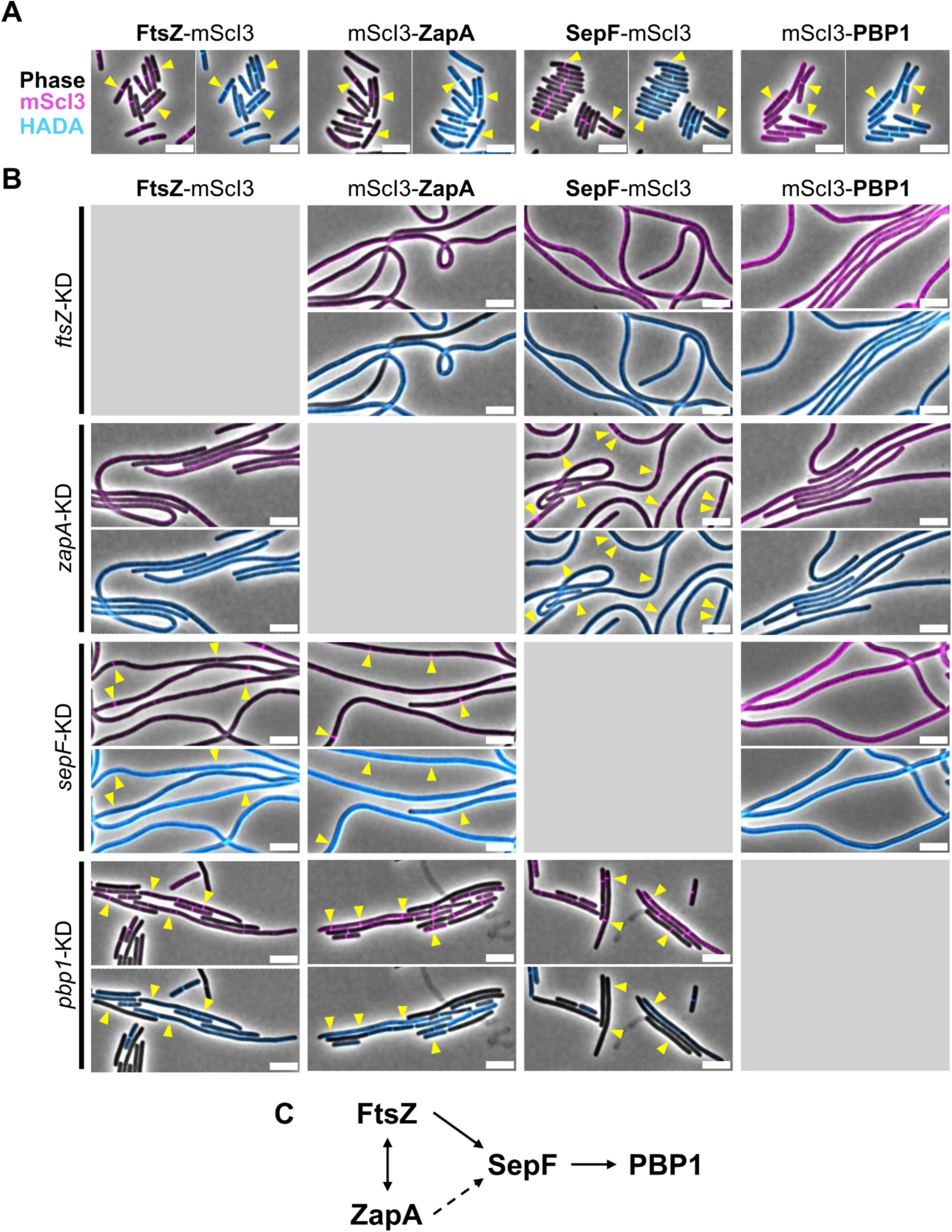
Order of assembly of essential divisome proteins. (A-B) Constructs encoding tagged divisome proteins were each expressed from an ectopic locus under the control of an aTc-inducible P*_tet_* promoter; xylose-inducible CRISPRi-KD constructs are encoded on a plasmid. Cells were cultured with 2.5% xylose for 6 hr and pulsed with aTc (FtsZ-mScI3 = 0.5 ng/mL; mScI3-ZapA = 0.5 ng/mL; SepF-mScI3 = 1 ng/mL; mScI3-PBP1 = 2.5 ng/mL). for the last hour of the treatment before labeling with HADA and fixing cells for microscopy. (A) Control *C. difficile* strains producing the indicated divisome protein-mScarlet-I3 (mScI3) fusion and harboring a negative control CRISPRi cassette with non-targeting sgRNA were visualized by microscopy to ensure that the CRISPRi cassette itself does not impact the protein localization. Yellow arrows point to divisome protein foci. (B) *C. difficile* cells producing divisome protein-mScI3 fusions upon the indicated CRISPRi *ftsZ*-KD, *zapA*-KD, *sepF*-KD, and *pbp1*-KD cassettes were visualized by microscopy. Yellow arrows point to divisome protein foci. The HADA signal is scaled equally across all images, but the mScI3 signal was scaled independently for optimal protein localization The data in A-B are representative of at least two independent experiments. Scale bars = 5 μm. (C) The order of assembly of the essential divisome proteins is depicted. The dotted line indicates that while SepF can form foci without ZapA, the foci appear irregular and diffuse, suggesting that ZapA is partially required for proper SepF localization.

We next analyzed the localization dependency of divisome proteins in the absence of SepF. We found that FtsZ-mScI3 and mScI3-ZapA could assemble into foci in *sepF*-KD cells, albeit at a lower frequency along the cell length than in control cells (**Fig. 4B**). These data indicate that SepF depends on FtsZ and partially on ZapA for assembly at mid-cell, but both FtsZ and ZapA can assemble in the absence of SepF. Thus, SepF is downstream of FtsZ and ZapA in the divisome assembly pathway (**Fig. 4C**). We also found that mScI3-PBP1 localized solely to the sidewall in the absence of SepF and failed to form the distinct foci observed in control cells (**Fig. 4B**). Therefore, SepF is required for PBP1 localization to the proto-ring complex, despite being dispensable for the assembly of the underlying FtsZ-ZapA complex. This loss of PBP1 localization likely explains why septum synthesis is blocked in the absence of SepF. Finally, we assessed the localization of divisome proteins during *pbp1*-KD. These analyses revealed that depletion of PBP1 does not prevent the assembly of FtsZ-mScI3, mScI3-ZapA, and SepF-mScI3 into foci, confirming that PBP1 recruitment occurs downstream of the proto-ring components FtsZ, ZapA, and SepF in the assembly pathway (**Fig. 4C**).

### Localization profile for non-essential divisome proteins MldA, MldB, MldC, DivIVA, FtsK, and PBP3

While these analyses revealed the localization dependencies of the essential divisome proteins FtsZ, ZapA, SepF, and PBP1, numerous other proteins have been shown to localize to the site of division that are not predicted to be essential in *C. difficile* (59, 63–65). Notably, non-essential genes may still play important roles during *C. difficile* cell division despite being dispensable for division in standard laboratory conditions for multiple reasons, including having a subtle regulatory function, being redundant with other genes, or exhibiting conditional essentiality. We therefore generated mScI3 fusions to several non-essential mid-cell localizing proteins, including MldA, MldB, MldC, DivIVA, FtsK, and PBP3.

MldA is single-pass transmembrane protein that carries an extracellular SPOR domain and a cytosolic globular domain and is encoded in an operon with MldB and MldC (64). These proteins are only encoded in close relatives of *C. difficile*, and their function remains unclear (64).

DivIVA is a late-stage division protein widely conserved across the Firmicutes, and it is known to localize to sites of negative membrane curvature (95, 104–106). *C. difficile* DivIVA has been shown to localize to septa (65), although heterologous expression in *B. subtilis* and biochemical interaction studies suggest that *C. difficile* DivIVA behaves differently from previously characterized systems (107, 108). Despite these analyses, DivIVA function in *C. difficile* has not been systematically examined.

FtsK is a single-pass transmembrane protein important for divisome assembly and chromosome segregation in many model systems (109–114). FtsK has also been localized to the division apparatus in *C. difficile* (59, 63), although it is not strictly required for *C. difficile* cell division (63, 66). Finally, PBP3 is a non-essential class B PBP that we recently found non-catalytically promotes the activity of PBP1 (62). While each of these proteins have been shown to localize to the site of division (59, 62–65), whether they localize to the mid-cell prior to, during, or after the onset of septum synthesis has not yet been determined, with the exception of PBP3, which we found localizes concurrently with septum synthesis (62).

To analyze the localization of MldA, MldB, MldC, DivIVA, FtsK, and PBP3 in *C. difficile*, we used a similar dilute-labeling approach where constructs encoding mScI3 fusions to each of these proteins were integrated into the genome under control of the aTc-inducible P*_tet_* promoter (**Fig. 5A**). Each fusion construct encodes either a GSAGSAAGSGKL linker (for MldA, MldB, and MldC) or (GGGGS)_3_ linker (for DivIVA, FtsK, and PBP3). To visualize the mid-cell localization for each of these non-essential mid-cell localizing proteins in relation to the onset of septum synthesis, we again generated demographs analyzing the localization of these proteins as a function of cell length (**Fig 5B**). We found that all six proteins appear to localize to the mid-cell later in the cell cycle than FtsZ-mScI3, mScI3-ZapA, or SepF-mScI3 (**Fig 2B**)(**Fig 5B**), with the localization of mScI3-MldB, mScI3-MldC, DivIVA-mScI3, FtsK-mScI3, and mScI3-PBP3 to mid-cell being largely coincident with the appearance of septa, as detected by HADA labeling (**Fig 5B**). In contrast, mScI3-MldA showed strong mid-cell localization prior to clear HADA labeling of septa (**Fig 5B**), suggesting that MldA localization to mid-cell occurs either prior to or early at the onset of septum synthesis.

**Fig 5:**
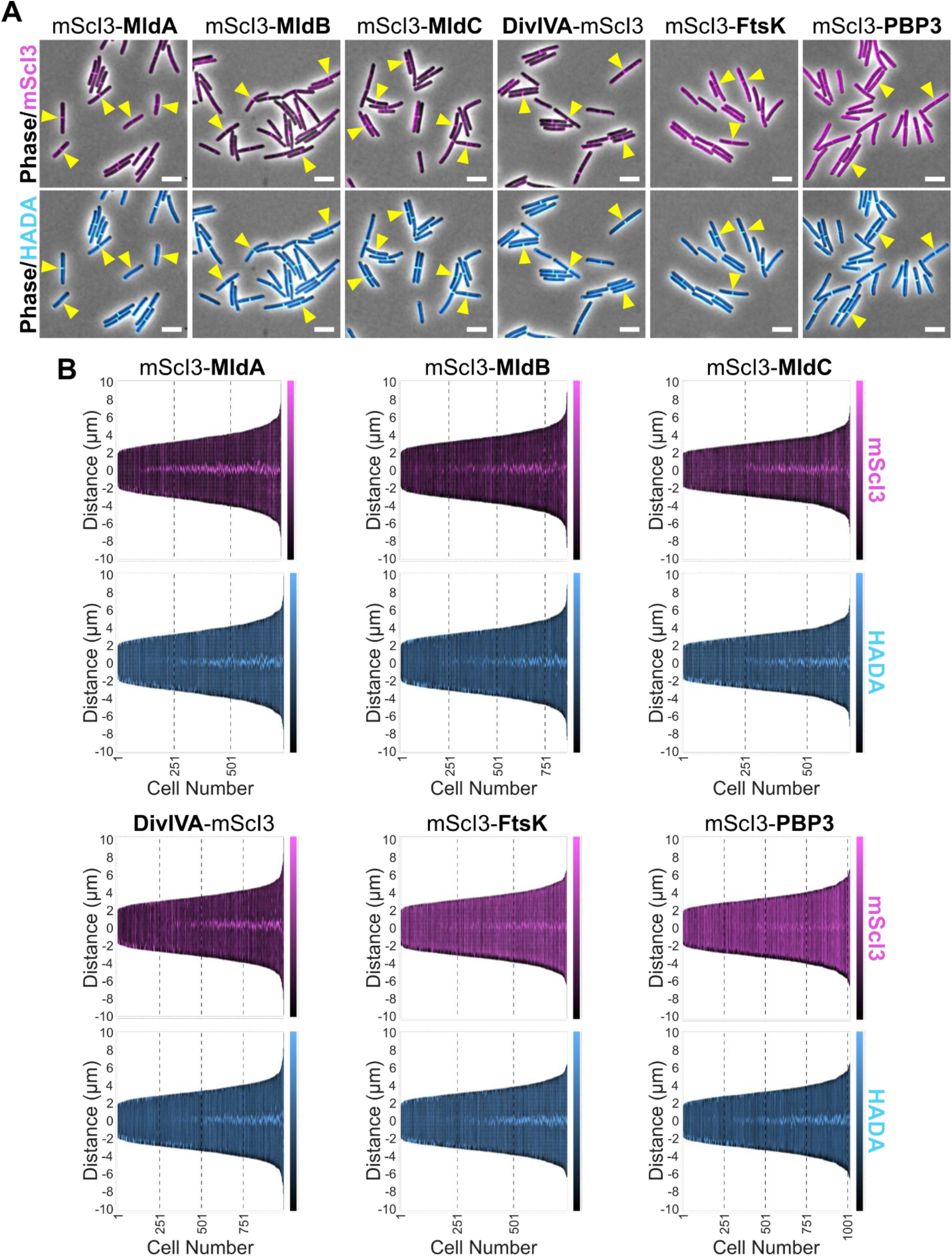
Localization profile of non-essential divisome proteins. (A-B) Constructs encoding mScarlet-I3 (mScI3) fusion proteins under the control of an aTc-inducible P*_tet_* promoter were expressed from an ectopic site in the genome. To induce production of the fusion protein, logarithmically growing cells were cultured in the presence of aTc for 1 hr (mScI3-MldA = 1 ng/mL; mScI3-MldB = 0.5 ng/mL; mScI3-MldC = 0.5 ng/mL; DivIVA-mScI3 = 0.5 ng/mL; mScI3-FtsK = 5 ng/mL; mScI3-PBP3 = 5 ng/mL). Cells were labeled with HADA for 10 min to visualize sites of peptidoglycan synthesis and/or remodeling and then fixed for fluorescence microscopy. (A) Merged images containing phase and mScI3 signal (top) or phase and HADA signal (bottom) are shown, and yellow arrows point to examples of divisome protein foci. Scale bars = 5 μm. (B) Demographs were generated using MicrobeJ to visualize the medial axis fluorescence profile of ≥500 cells. Cells are ordered by length, and the mScI3 or HADA signal along the distance of the cell relative to mid-cell is shown in magenta and cyan, respectively. Data in A-B are representative of at least three independent experiments.

### Dependence of MldA, MldB, MldC, DivIVA, FtsK, and PBP3 on FtsZ, SepF, and PBP1 for mid-cell localization

Since the four, core essential divisome proteins are recruited stepwise in a hierarchical order consisting of (i) FtsZ and ZapA, (ii) SepF, and (iii) PBP1 (**Fig 4C**), we next sought to identify the localization dependencies for the non-essential divisome proteins. To this end, we localized fluorescent protein fusions of the non-essential divisome proteins upon KD of *ftsZ*, *sepF*, or *pbp1*, which represent the three stages of *C. difficile* divisome complex assembly. We found that none of the non-essential divisome proteins localized in *ftsZ*-KD cells, and upon *sepF-KD*, no foci were observed for mScI3-MldA, mScI3-MldC, DivIVA-mScI3, mScI3-FtsK, and mScI3-PBP3 (**Fig 6A**). Intriguingly, mScI3-MldB formed foci in *sepF*-KD cells, suggesting that MldB depends on FtsZ but not SepF for localization.

**Fig 6:**
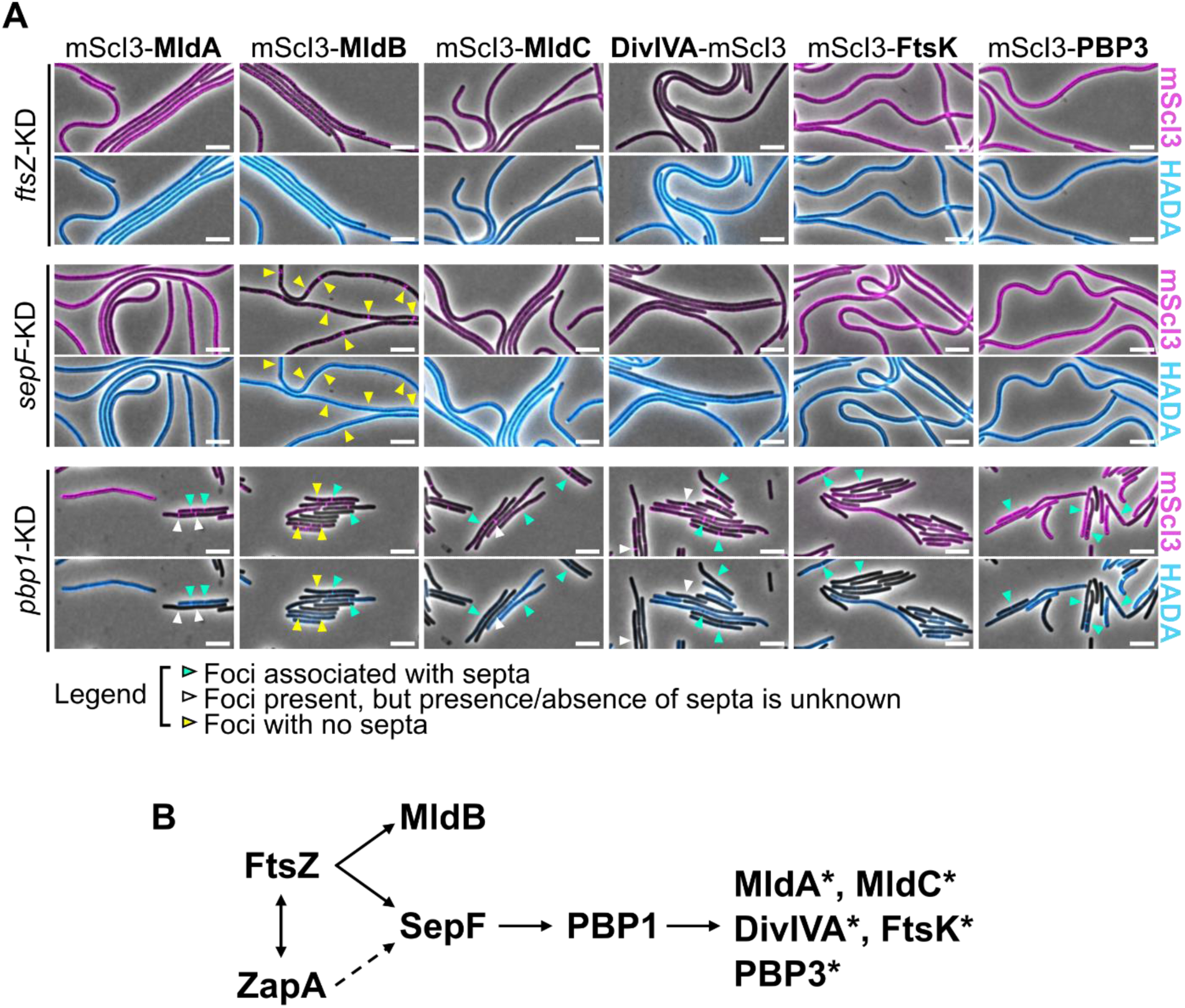
Dependence of non-essential divisome proteins on FtsZ, SepF, and PBP1 for localization. (A) Constructs encoding fluorescent protein fusions to divisome proteins were expressed from an ectopic locus under the control of an aTc-inducible P*_tet_* promoter during xylose-inducible CRISPRi-KD of the indicated genes. The CRISPRi-KD constructs were expressed from a plasmid. Cells were cultured with 2.5% xylose for 6 hr and pulsed with aTc for the last hour of the treatment (mScI3-MldA = 1 ng/mL; mScI3-MldB = 0.5 ng/mL; mScI3-MldC = 0.5 ng/mL; DivIVA-mScI3 = 0.5 ng/mL; mScI3-FtsK = 5 ng/mL; mScI3-PBP3 = 5 ng/mL) before labeling with HADA and fixing cells for microscopy. Divisome protein foci are indicated by arrows. Cyan arrows represent divisome protein foci that are co-localized with septa in the *pbp1*-KD strain, which is likely due to incomplete depletion of PBP1 protein leading to a low-level of septum synthesis. White arrows represent divisome protein foci within cells that did not label efficiently with HADA, so the presence or absence of septa at the site of protein localization is unknown. Yellow arrows indicate divisome protein foci that localize in the absence of septa, specifically in cells that label well with HADA. Data are representative of at least two independent experiments. Scale bars = 5 μm. (B) The order of divisome assembly is depicted. The dotted line indicates that ZapA is only partially required for proper SepF localization. The proteins labeled with an * indicate that the protein only formed foci when septa were detectable; these septa likely form because PBP1 is only partially depleted.

The localization hierarchy was more difficult to assess during *pbp1* KD because some *pbp1*-KD cells form septa with associated divisome protein foci. We reasoned that septal co-colocalization was most likely due to incomplete depletion of PBP1, since only ∼75% PBP1 is depleted during the CRISPRi KD based on prior results (62). To determine if a protein was capable of localizing independently of PBP1, we therefore looked specifically for divisome protein foci that did not co-localize with septa. It should be noted that this approach was further complicated by the heterogeneity in HADA labeling observed during *pbp1* KD, with a proportion of *pbp1*-KD cells failing to label with HADA entirely (**Fig 6A**)(62); thus, the presence or absence of septa in some cells could not be determined. Despite these limitations, we found that mScI3-MldA, mScI3-MldC, DivIVA-mScI3, mScI3-FtsK, and mScI3-PBP3 all formed mid-cell foci in the *pbp1*-KD cells that were either co-localized with septa or in cells that did not label with HADA (**Fig 6A**). Only mScI3-MldB could form foci that were not clearly associated with septa (yellow arrows, **Fig 6A**), consistent with MldB not being dependent on PBP1 for recruitment to the divisome complex. Collectively, our data support that MldA, MldC, DivIVA, FtsK, and PBP3 are likely dependent on the presence of FtsZ, SepF, and PBP1 to be recruited to the divisome, whereas MldB is dependent only on FtsZ (**Fig 6B**). Thus, with the exception of MldB, the non-essential divisome proteins MldA, MldC, DivIVA, FtsK, and PBP3 appear to depend on septum synthesis to localize to mid-cell.

To more precisely analyze the role of these non-essential proteins during cell division, we generated clean, in-frame deletion mutants for each of the genes encoding non-essential divisome proteins analyzed above. This genetic strategy allowed us to avoid the potentially off-target or polar effects caused by the genetic strategies previously used to study many of these non-essential divisome genes, including Targetron disruption (MldA, MldB, or MldC) (64), antisense RNA (FtsK) (59), or transposon-insertion (63, 66).

The Δ*mldA*, Δ*mldB*, Δ*mldC*, Δ*divIVA*, Δ*ftsK*, and Δ*pbp3* mutants all completed cell division and form septa (**Fig 7A**, yellow arrows), consistent with these genes being dispensable for division. However, we found that clean deletion of Δ*mldA*, Δ*mldB*, and Δ*mldC* led to a mild chaining phenotype (**Fig 7A**), similar to prior Targetron mutants targeting *mldA* and *mldB* (64). Importantly, we could complement the Δ*mldA* mutant and reverse its chaining phenotype by expressing *mldA* from an ectopic site in the genome (**Fig 7B**). The Δ*divIVA* deletion mutant also grew as short chains (**Fig 7A**), phenocopying the CRISPRi *divIVA*-KD mutant (**Fig 1C**). The chaining phenotype could be complemented by expressing *divIVA* under the control of the constitutive P*_cwp2_* promoter from an ectopic site. Intriguingly, although an *ftsK*-antisense KD mutant had previously been shown to exhibit a filamentation phenotype (59), we found that a Δ*ftsK* clean deletion mutant had a WT morphology (**Fig 7A**). Together, our data support that these non-essential divisome proteins are not required for septum synthesis and likely play an accessory and/or regulatory role in *C. difficile* division.

**Fig 7:**
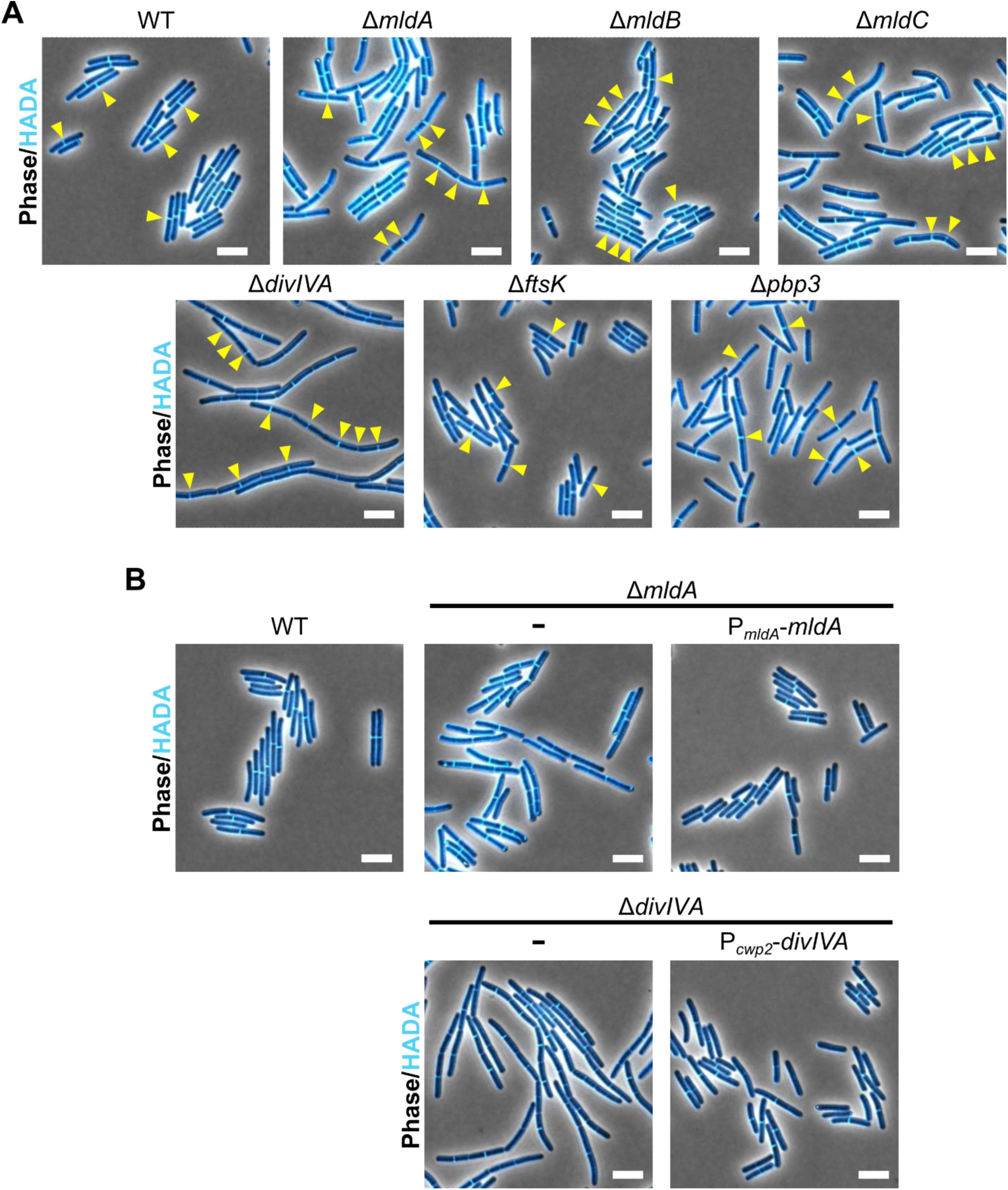
Phenotypes of deletion mutants lacking non-essential divisome genes. (A-B) The indicated clean deletion strains of *C. difficile* were cultured to logarithmic phase and labeled with HADA. (A) yellow arrows point to division septa. (B) The Δ*mldA* or Δ*divIVA* mutants were complemented with either an empty vector integrated into an ectopic locus “-“ or the indicated expression construct. Scale bars = 5 μm.

## Discussion

Despite the highly conserved nature of cell division in bacteria (39), the majority of canonical divisome proteins characterized in bacteria are either missing from *C. difficile*’s genome (FtsW, FtsI, FtsA, EzrA, GpsB) or their function has not been preserved during vegetative cell division (FtsQ, FtsL, FtsB). Since this essential process remained poorly characterized in *C. difficile*, in this study, we gained insight into how *C. difficile* assembles its divisome by determining the order of assembly of four essential divisome proteins FtsZ, ZapA, SepF, and PBP1, along with 6 non-essential mid-cell localizing proteins. We also genetically defined the specific requirement for SepF and show that several non-essential mid-cell localization proteins are dispensable for septum synthesis.

These analyses revealed that ZapA plays a critical role during *C. difficile* FtsZ-ring formation (**Fig 4**), in contrast with most model systems studied to date, where ZapA is dispensable for this process due to the presence of multiple redundant FtsZ-bundling mechanisms (4, 71–77). Similarly, while SepF is often dispensable in Firmicutes species due to the presence of additional FtsZ membrane tethers (5, 6, 78, 115), we found that SepF is essential for septum synthesis in *C. difficile* (**Fig 1**). Our studies further revealed that FtsZ/ZapA rings form in the absence of SepF in *C. difficile* (**Fig 4**). Since the septum synthase PBP1 depends on SepF to be recruited to marked division sites (**Fig 4**), these analyses suggest that the primary function of SepF in *C. difficile* is to link the cytoskeletal FtsZ/ZapA scaffold to late-stage divisome proteins, including PBP1. Thus, rather than promoting FtsZ-ring stability or membrane tethering, SepF may play an important role in licensing the progression of cell division. The localization dependence of SepF on FtsZ is similar to other Firmicutes such as *S. suis* and *B. subtilis*, where SepF localization depends on FtsZ, but FtsZ can still assemble without SepF (Hamoen 2006 *Mol Micro*)(Gao 2025 BMC Microl). These findings are also reminiscent of the localization dependency observed in archaeal species that encode SepF as the ancestral FtsZ membrane anchor, where SepF depends on FtsZ1 to assemble at mid-cell, but FtsZ1 and FtsZ2 can still assemble into rings when SepF is depleted, at least in *Haloferax volcani* (83). Notably, in addition to its proposed FtsZ membrane-anchoring function, SepF functions as an interaction hub for divisome proteins in several organisms. In *C. glutamicum*, SepF assembles into a complex with the FtsZ-interacting gephyrin-like protein Glp and its transmembrane receptor GlpR (116), and in Archaea, SepF organizes the photosynthesis reaction center proteins CdpB1/2/3, which are important for division in Archaea (117, 118), and the archaeal cell division protein CdpA (119). Thus, understanding how *C. difficile* SepF contributes to the recruitment of late divisome proteins, such as PBP1, should advance our understanding of how this pathogen assembles a functional divisome.

Our work also provides insight into the recruitment and function of the non-essential mid-cell localizing proteins MldA, MldB, MldC, DivIVA, FtsK, and PBP3. While most of these proteins localize to mid-cell coincident with septum synthesis, MldB can localize to mid-cell independent of septum synthesis, including in the absence of SepF and PBP1 (**Fig 6**). Since our clean deletion mutant analyses showed that Δ*mldA*, Δ*mldB*, Δ*mldC*, and Δ*divIVA* mutants all exhibit a chaining phenotype, presumably due to a defect in cell separation after septum synthesis is complete (**Fig 7**), we propose that most of these non-essential proteins play a role in late-stage cell division. These genetic analyses build on prior work using Targetron insertions and plasmid-based complementation to individually analyze MldA, MldB, and MldC function during cell division, reinforcing the previous conclusion that each of these genes is required for proper cell separation (64)(**Fig 7A**). Our analyses also establish a role for DivIVA during *C. difficile* cell separation. While the mechanism by which DivIVA regulates this process in *C. difficile* remains unclear, in some Firmicutes, DivIVA appears to promote cell separation often by regulating the export of cell separation enzymes (93–97, 120). Interestingly, our clean deletion analyses showed that deletion of *ftsK* does not result in a cell division defect (**Fig 7**), in contrast with prior work indicating that antisense RNA-based KD of *ftsK* causes filamentation (59). Deletion of *ftsK* may produce a different phenotype from antisense RNA KD due to compensatory transcriptional changes in the deletion mutant, off-target or polar effects of the antisense RNA, or differences in culture conditions or CD630Δ*erm* isolates between labs.

Our analyses provide further insight into how the orphan Class B PBP enzyme PBP3 modulates both vegetative cell division and asymmetric division during sporulation (62, 121). We previously implicated PBP3 in promoting PBP1 glycosyltransferase activity during vegetative cell division (62) and asymmetric division (121). Since our prior work indicated that in both cases PBP3 function does not depend on its own catalytic activity, PBP3 likely regulates *C. difficile* cell division via protein-protein interactions with the divisome complex (62, 121). We build upon these prior findings by demonstrating that PBP3 localization to the site of division depends on FtsZ, SepF, and PBP1 (**Fig 6**). It should be noted that significant processing of the mScI3-PBP3 fusion protein limits the sensitivity of these localization assays. Still, precisely how PBP3 regulates the *C. difficile* divisome remains to be determined.

Finally, our study optimizes CRISPRi-compatible trans-complementation methods for rapidly testing the function of FtsZ, ZapA, and SepF variants in *C. difficile*. Indeed, our understanding of FtsZ, ZapA, and SepF function in *C. difficile* lags behind model systems, where dozens of mutant variants have been evaluated to dissect the function of these proteins at the molecular level. To our knowledge, not a single point mutant has yet been evaluated in *C. difficile* for FtsZ, ZapA, or SepF. Our newly established conditional expression system will facilitate rapid structure-function analyses of these core divisome proteins in the future, bypassing more laborious strategies for generating chromosomally-encoded conditional expression strains. Indeed, our trans-complementation system allowed the functionality of tagged divisome proteins and their potential processing to be directly addressed. These analyses revealed that an FtsZ-mScI3 fusion is inactive in *C. difficile* when expressed as the sole copy of FtsZ, whereas mScI3-ZapA, SepF-mScI3, and mScI3-PBP1 each have at least partial activity (**Fig 3**). It should be noted that it has been challenging to identify functional fusions to FtsZ, with N- and C-terminal FtsZ-fluorescent protein fusions either being non-functional or causing suppressor mutations in *E. coli* (99, 122), or requiring low temperatures for function, as in *B. subtilis* (8). Furthermore, for *E. coli* FtsZ, it took nearly 20 years before functional sandwich fluorescent protein fusions to FtsZ were identified, and still these fusions cause subtle abnormalities at 37°C (123). Since fluorescent protein fusions to the C-terminus of FtsZ are functional in *Streptococcus pneumoniae* (93, 124), exploring different C-terminal tags other than mScI3 or implementing a sandwich fusion strategy for *C. difficile* FtsZ may lead to functional fusions in the future.

Altogether, our work supports a core function for FtsZ, ZapA, SepF, and PBP1 in *C. difficile* cell division, and supports prior work showing that the *C. difficile* divisome differs substantially from model systems (41, 63). In addition to the genes analyzed in this study, there are likely unidentified divisome proteins important for this process. For example, how PBP1 is recruited to the divisome complex and how its activity is licensed at mid-cell to drive septum synthesis and cytokinesis remains to be determined. The methods for conditional gene expression and protein localization optimized here will facilitate future mechanistic analyses of this unique and essential pathway.

## Materials and Methods

### Bacterial strains and growth conditions

*C. difficile* strains are listed in Table S1 and derived from the 630Δ*erm* strain background. Chromosomally-encoded mutations, including insertion of *mScI3*-tagged constructs and gene deletions were generated in a Δ*pyrE* strain using *pyrE*-based allele coupled exchange as previously described (125). *C. difficile* strains were cultured in brain heart infusion medium supplemented with 0.5% yeast extract and 0.1% L-cysteine (BHIS) with thiamphenicol (10 μg/mL) to maintain plasmids, and/or kanamycin (50 μg/mL) and cefoxitin (8 μg/mL), and/or spectinomycin (500 μg/mL) as needed for genetic manipulation. *C. difficile* defined medium (CDDM) (126) was used to select for *pyrE* restoration during allele-coupled exchange, or was supplemented with 5-fluoroorotic acid (2 mg/mL) and uracil (5 μg/mL) to counter-select against *pyrE* for making clean deletion mutants, with spectinomycin (500 μg/mL) included to select for the *aad9* marker when making a marked deletion (e.g. Δ*ftsK::aad9*). Deletion mutants were verified by PCR with locus-specific primers. *C. difficile* cultures were grown at 37°C in an anaerobic chamber containing a gas mixture comprised of 85% N_2_, 5% CO_2_, and 10% H_2_.

Plasmids used in this study were cloned by Gibson assembly, propagated in *E. coli* DH5α, and sequence-confirmed by Sanger or nanopore sequencing. *E. coli* strains and plasmids used in this study are listed in Table S2, with links to plasmid maps that include the primer sequences used for cloning. To introduce constructs into *C. difficile*, plasmids were first transformed into *E. coli* HB101/pRK24 and then conjugated into C. difficile as previously described (127). *E. coli* cultures were grown in LB at 37°C supplemented as needed with chloramphenicol (20 μg/mL), ampicillin (100 μg/mL), or kanamycin (30 μg/mL).

### CRISPRi-KD and CRISPRi-compatible complementation

A plasmid-encoded CRISPRi-KD system based on the pIA33 plasmid was used as previously described (67). Briefly, sgRNAs were designed to target the non-targeting strand, and were chosen using the Benchling sgRNA design tool. These were cloned downstream of the constitutive P*_gluD_* promoter. Plasmids were maintained in *C. difficile* by selection with 10 μg/mL chloramphenicol. CRISPRi-KD was induced by culturing strains in the presence of 2.5% xylose, back-diluting as needed to maintain logarithmic growth in the presence of inducer for 5-6 hr. When conditional expression of CRISPRi-resistant complementation constructs was required, constructs containing synonymous mutations in the sgRNA-targeted sequence were integrated into the genome downstream of the *pyrE* locus under the control of an aTc-inducible P*_tet_* promoter. Cultures were maintained in the presence of both xylose and aTc to conditionally express the CRISPRi-resistant complementation construct while knocking down expression of the endogenous gene.

### Fluorescent probes

The fluorescent D-amino acid HADA (Tocris) was used to label sites of peptidoglycan synthesis and/or remodeling. 500 μL of logarithmically growing *C. difficile* was exposed to 50 μM HADA for 10 min, then fixed with a mixture of 100 μL 16% paraformaldehyde and 20 μL 1 M NaPO_4_ buffer (pH 7.4) for 30 min at room temperature and 30 min on ice (61). Fixed cells were washed three times with 1 mL 1XPBS prior to imaging.

### Protein localization with mScarlet-I3 fusions

*C. difficile* strains were engineered to express chromosomally-encoded divisome protein-mScI3 fusions under control of the aTc-inducible P*_tet_* promoter, integrated downstream of *pyrE*. Strains were cultured to the logarithmic phase in BHIS and then exposed to the indicated concentration of aTc for 1 hr. Induced cultures were labeled with HADA and fixed as described above. After cell fixation, cells were incubated overnight at room temperature in the dark to allow chromophore maturation, as previously described (61).

### Fluorescence microscopy

Microscopy samples were imaged on agarose pads (1% agarose in 1XPBS). Phase-contrast and fluorescence micrographs were acquired with a Leica DMi8 inverted microscope equipped with a 63X 1.4 NA Plan Apochromat oil-immersion phase-contrast objective, a high precision motorized stage (Pecon), and in a 37°C incubator (Pecon). Excitation light was generated by a Lumencor Spectra-X multi-LED light source with integrated excitation filters. An XLED-QP quadruple-band dichroic beam-splitter (Leica) was used (transmission: 415, 470, 570, and 660 nm) with an external filter wheel for all fluorescent channels. HADA was excited at 395/25, and emitted light was filtered using a 440/40-nm emission filter, with a 120 ms exposure time; mScarlet-I3 was excited at 550/28 nm, and emitted light was filtered using a 590/50-nm emission filter, with a 150 ms exposure time. Light was detected using a Leica DFC 9000 GTC sCMOS camera. 1–2 μm z-stacks were taken with 0.21 μm z-slices. Images were acquired using the LASX software, and fluorescence images were deconvolved using Leica Small Volume Computational Clearing with the following settings: refractive index 1.33, strength 60%, and regularization 0.05.

Images were processed using FIJI to select the best-focused z-plane for each channel and adjust the image brightness and contrast of images. Identical minimum and maximum values were applied to a given channel across all images in a figure panel so that direct comparisons could be made across the images within a figure panel, unless otherwise specified in the figure legend. Demographs were generated with MicrobeJ in FIJI, using segmentation masks generated by SuperSegger (128, 129).

### Western blot analysis

A volume of 1.4-2.8 mL of logarithmically growing *C. difficile* culture (OD ∼0.4-.07) were pelleted, resuspended in 25 μL 1X PBS, freeze-thawed three times, mixed with 25 μL EBB buffer (9 M urea, 2 M thiourea, 4% SDS, 2 mM β-mercaptoethanol), and boiled for 20 min to lyse cells. SDS-polyacrylamide gel electrophoresis (SDS-PAGE) was performed on 10% polyacrylamide gels. Proteins were transferred to polyvinylidene difluoride membranes, which were subsequently probed with the following primary antibodies: rabbit polyclonal anti-DivIVA TF142 (62) at 1:1,000 dilution; chicken polyclonal anti-GDH (aCdGDH; Thermo) at 1:10,000 dilution; and/or a goat polyclonal anti-mScarlet at 1:5,000 dilution (MBS448290; MyBioSource). Anti-rabbit, anti-chicken, and anti-goat IR800 or IR680 secondary antibodies (LI-COR Biosciences, 1:30,000) were used to detect bands with a LI-COR Odyssey CLx imaging system.

## Acknowledgements

This work was supported by the National Institute of Allergy and Infectious Diseases grant R01 AI122232 to AS. Additionally, GAH was supported by the Tufts University Institutional Research Career and Academic Development Award Program (K12GM133314) and a National Institute of Allergy and Infectious Diseases training fellowship (F32AI191529). The content is solely the responsibility of the authors and does not necessarily represent the official views of the National Institutes of Health. The funders had no role in study design, data collection and analysis, decision to publish, or preparation of the manuscript.

